# Oxygen changes drive non-uniform scaling in *Drosophila melanogaster* embryogenesis

**DOI:** 10.1101/024208

**Authors:** Steven G. Kuntz, Michael B. Eisen

## Abstract

In *Drosophila* embryogenesis, increasing either oxygen concentration or temperature accelerates development. Having previously investigated temperature’s impact on embryogenesis, we characterized developmental response to oxygen levels using time-lapse imaging. Changing oxygen concentrations greatly impact survival, with developmental rate changes that are dwarfed by those induced by temperature. While extreme temperatures increase early embryo mortality, mild hypoxia increases arrest and death during mid-embryogenesis and mild hyperoxia increases survival over normoxia. Though not independent, the reactions to temperature and oxygen are fundamentally different, with developmental time being inversely proportional to oxygen concentration but logarithmically related to temperature. Most notably, while development scales uniformly with temperature, oxygen changes drive developmental heterochrony. Morphological processes change with oxygen concentration at different rates. Gut formation is more severely slowed by decreases in oxygen, while head involution and syncytial development are less impacted than the rest of development. These data reveal that uniform scaling, seen with changes in temperature, is not the default result of adjusting developmental rate.

## Introduction

Vital for respiratory energy production, oxygen plays a central regulatory role in growth and metabolism. The basic molecular mechanism of oxygen sensing has been worked out [1–3], but its impact on individual morphological processes remain unknown. *Drosophila* frequently lay eggs on substrates colonized by bacteria and yeast, which affect local oxygen concentrations both in the lab and in the field. With microbial overgrowth, oxygen levels can fall precipitously.

Hypoxia tolerance varies between tissues [2] and possibly between stages of embryonic development. Either cellular growth or migration may affect oxygen consumption, making some stages more resistant to arrest. Hypoxic *Drosophila* syncytial embryos readily arrest at a metaphase checkpoint [4] and resume development under normoxia if the hypoxic period is not too long [5]. Cellularized embryos survive longer hypoxic periods, up to several days [6]. Hypoxic arrest may not be entirely benign, as even brief periods of hypoxia lead to smaller bodies and wings, driven in part by decreased cell size [7]. Active oxygen sensing and nitric oxide signaling drive this arrest, which cannot be replicated using cyanide since it is independent of the electron transport chain [6, 8, 9].

*Drosophila*, like other animals, regulate metabolism and gene expression in response to changes in oxygen levels through the *HIF-1α* pathway, which communicates with the Tor and VEGF pathways. Under normal conditions, proline residues of *simalar* (*sima/hif-1/HIF-1α*) are hydroxylated by prolyl hydroxylase (*Hph/egl-9/EGLN*) to both inactivate *sima/hif-1/HIF-1α* and target it for Vhl-dependent degradation [1, 2]. The prolyl hydroxylase *Hph/egl-9* is itself negatively regulated under hypoxia by the cystathionine *β*-synthase *Cbs/cysl-1/CBS*, an ambient oxygen sensor via hyrodgen sulfide signaling. a*sima* has an oxygen-dependent degradation domain with a nuclear export sequence [3]. Thus, only during hypoxia does *sima* escape degradation and accumulate in the nucleus [1]. Rather than serving as a switch, the process is dynamic, with greater levels of oxygen accelerating both the degradation and nuclear export of *sima* [3].

The nature of the response to oxygen could be limited by transcription, enzyme kinetics, molecular processes, metabolism, and temperature. Transcriptional state changes significantly across the maternal-zygotic transition. Prolyl hydroxylation and Vhl-dependent degradation may influence the response. Nuclear import and export rates or intercellular signaling, since the HIF-1α pathway can be activated by insulin [10] in later stages, may affect the oxygen response. Metabolism has a non-linear effect on adult flies [11] and embryogenesis involves numerous metabolic changes [12]. All of these factors will be affected by ambient temperature, which also changes oxygen diffusion and solubility, though under normal conditions oxygen is not limiting even at high temperatures [13].

This work uses time-lapse imaging of embryos under a range of oxygen concentrations with precise temperature control to monitor the effects on developmental timing and morphology. In covering hypoxic through hyperoxic and warm through cold conditions, we have collected dynamic data on how the developing embryo responds to oxygen.

## Results

### Oxygen concentration controls developmental rate

We used automated time-lapse imaging in an airtight box with oxygen concentration control (*±*1%) and precise temperature control (*±*0.1°C) to track development using previously described methods [14]. We investigated embryos raised at constant oxygen concentrations (29%, 25%, 21%, 17%, 14%, and 10% *O*_2_) and kept at three different temperatures (17.5°C, 22.5°C, and 27.5°C), giving a total of eighteen specific conditions with over 800 embryos. A schematic of the setup is provided in Figure 1A. The actual setup is shown in Figure 1B.

**Figure 1.**
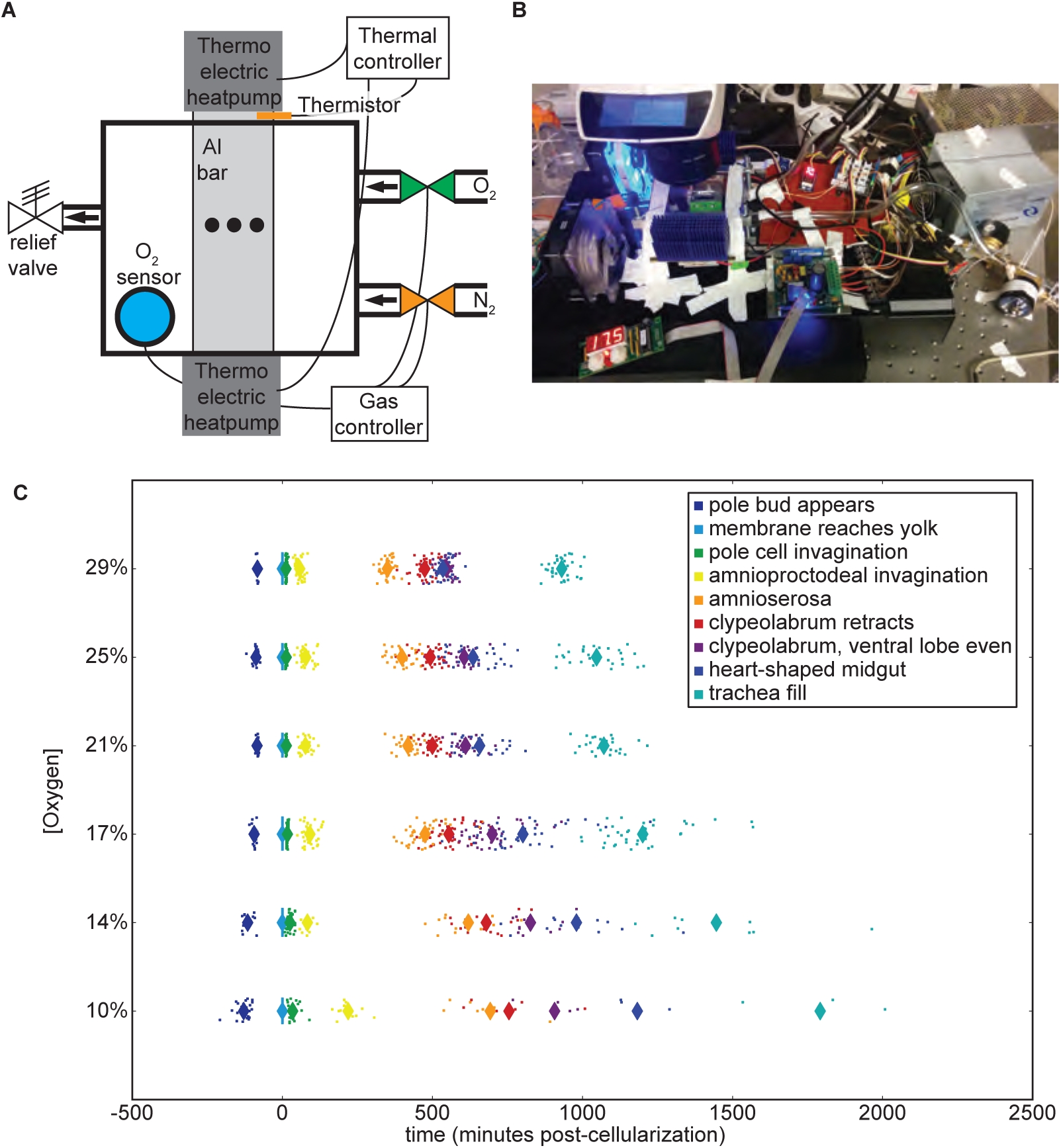
Developmental rate responds to oxygen concentrations. (A) The oxygen control schematic. A thermistor embedded in the aluminum bar provides temperature data to the temperature controller, which in turn adjusts the voltage to the thermo-electric controllers (Peltier). An oxygen sensor in the airtight box provides feedback on oxygen concentrations to the gas controller, which opens and closes oxygen and nitrogen valves accordingly. Embryos are imaged in the center of the aluminum bar within the airtight box, indicated by the black dots in the schematic. (B) Image of the oxygen control setup mounted on the microscope at 20% oxygen and 17.5°C. (C) Developmental rate across all stages changes with the oxygen concentration, performed at 27.5°C. Each animal is represented with a dot, with averages represented with a large diamond. Developmental times here are zeroed on the end of cellularization.

In agreement with previous research, developmental rate correlates with oxygen concentrations (Figure 1C). Hyperoxia accelerates development, allowing embryos to hatch sooner than they would under normal atmospheric conditions. Hypoxia slows development in a dose-dependent fashion. As oxygen levels fall, an increasing fraction of embryos die or arrest their development. Therefore, there are fewer embryos shown in Figure 1C at lower oxygen concentrations due to low rates of successful development, despite similar numbers of animals being prepared for imaging (Table S1).

### Timing of death and arrest depend on both oxygen and temperature

Oxygen levels affect the stage at which embryos arrest or die. Higher concentrations of oxygen (29%) lead to more animals dying during early development, including death in the syncytium and a failure to properly gastrulate. This point of failure is similar to that observed at high temperatures with normal oxygen levels [14]. Lethality at 25% oxygen is actually lower than that at 21%, which approximates atmospheric levels. Problems with development may be aggravated by the de-chorionation and mounting procedure. At high temperatures (32.5°C) and high oxygen (29%), almost all embryos die very early in development (Table S1).

At lower oxygen levels there is a major shift from very early developmental arrest and death to mid-embryogenesis arrest (Figure 2). This holds true at all temperatures (especially at 10% *O*_2_), but is most pronounced at 27.5°C, where the effects are still seen at 14% *O*_2_. Frequently development halts during germ-band retraction, preventing full exposure of the amnioserosa. The midgut primordia in these embryos routinely migrates haphazardly after arrest, coinciding with the embryo falling into morphological disarray. In embryos that pass these mid-embryogenesis stages, trachea formation often proves problematic. Commonly the trachea fails to form, which coincides with arrest late in midgut formation, following the heart-shaped midgut stage. These animals generally form functional muscle, with some twitching observed.

**Figure 2.**
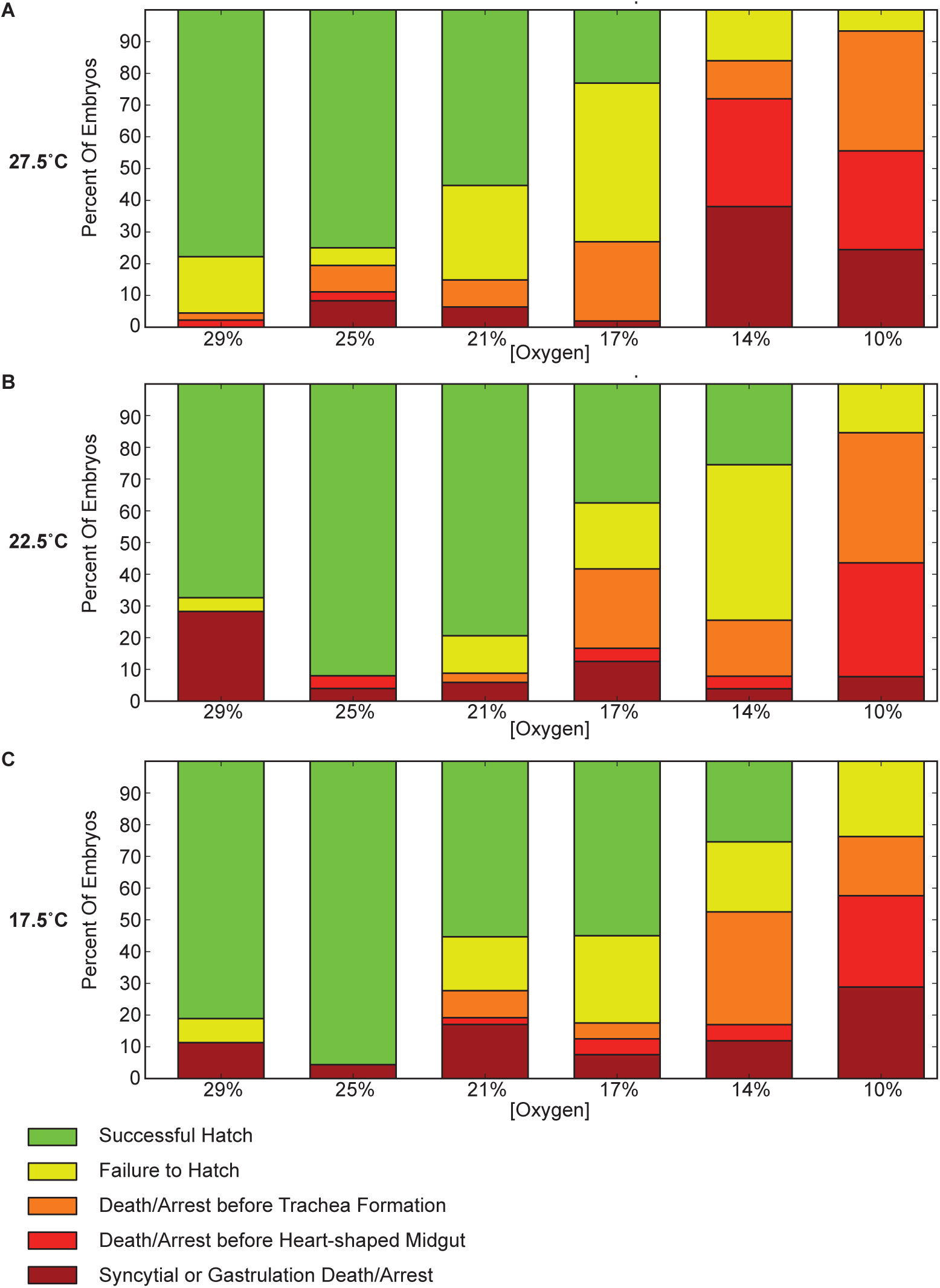
The point of failure in development depends on both oxygen and temperature. Lethality at different concentrations are shown for three different temperatures (27.5°C, 22.5°C, and 17.5°C). Lower concentrations of oxygen are more likely to exhibit failure of during pre-tracheal development, with a particularly large increase in mortality between gastrulation and completion of the heart-shaped midgut (shown in red). A substantial increase in late development before trachea fill is also seen (shown in orange). Developmental arrest is frequently at germ band retraction. This is in contrast with higher oxygen concentrations, where failure is almost exclusively very early in development (shown in brick red), prior to the completion of gastrulation, or during difficulties hatching following trachea filling (shown in yellow). Highest survival is interestingly at 25% oxygen.

Higher, but still hypoxic, oxygen levels (14% and 17%) have a significant fraction of embryos that fail to hatch. While embryonic development appears to be completed, including the filling of the trachea with air, larvae struggle to break out of their vitelline membrane yet fail to escape. While seen in all conditions, this behavior is most prevalent in these mildly hypoxic conditions.

### Temperature influences oxygen’s control of developmental rate

Decreasing oxygen concentrations from 29% to 10% at any temperature lead to an additional sixteen to eighteen hours of developmental time (Figure 3A). This results in a different proportional change at each temperature, with nearly a 100% increase at 27.5°C and only a 50% increase at 17.5°C. This contrasts with changes in temperature, where developmental time roughly doubles over a 10°C range, regardless of the oxygen concentration (Supplementary Figure S1).

**Figure 3.**
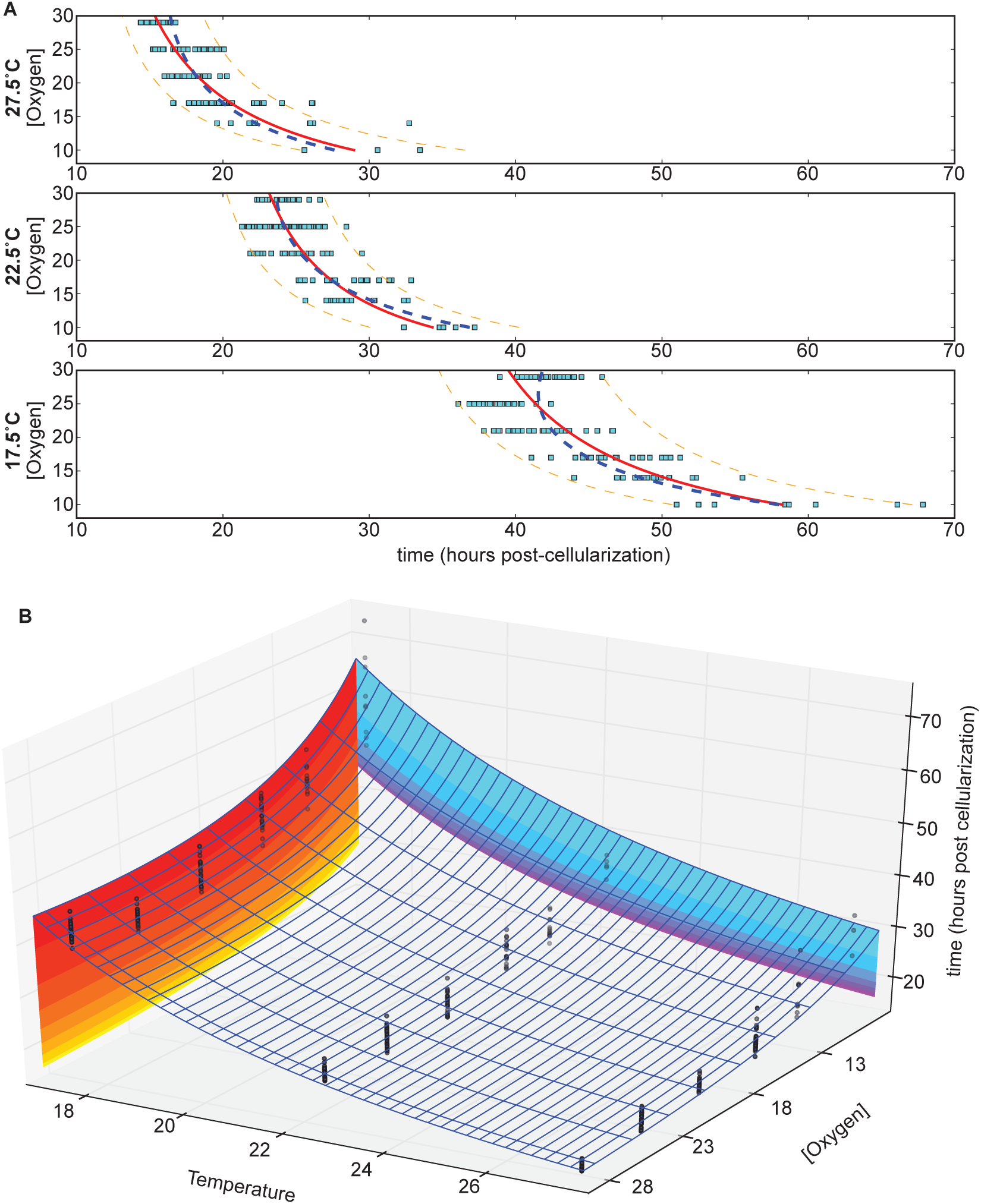
Oxygen and temperature affect developmental time cooperatively. (A) Fitting at different temperatures (27.5°C, 22.5°C, 17.5°C). The shift is very temperature dependent. The solid red line represents a fit at each particular temperature, with 90% confidence of reproduction marked with the dashed orange line. The blue dashed line represents the fit across all temperatures, which departs from the individual temperature fits. (B) Total developmental time is affected by both temperature and oxygen levels. Each point represents an individual embryo at a given temperature and oxygen level. Color contours help visualize the transitions of increased heat (yellow to red contour) and increased oxygen (purple to blue contour).

Changes in oxygen concentration have an inverse proportional effect on developmental rate. Least squares curve fitting was attempted with multiple models, including exponential models, for changes in oxygen concentration. The data most closely matched a Monod equation model. The parameters of the response change significantly with temperature, however the qualitative response is the same (Figure 3A):

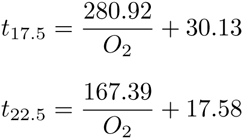

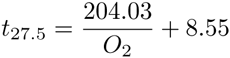

Fitting at each oxygen concentration (Supplementary Figure S1) yields relatively good fits using an exponential Arrhenius model. These different methods of fitting can be combined and yield the best fit as a multivariable non-additive model. The overall effect of oxygen and temperature can be combined to yield (Figure 3B):

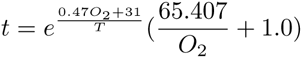

Based on the fit, oxygen appears to have an effect on both the linear and exponential coefficients. This model is empirical and does not predict effective oxygen concentration as a function of temperature-dependent changes in oxygen solubility and diffusion. Increased oxygen may allow some additional growth acceleration, but the acceleration of growth rate appears to be leveling off, asymptotically approaching a maximum. At lower oxygen levels, the prevalence of arrest is expected to overtake the observed response curve.

### Oxygen dependent developmental scaling is non-uniform and temperature-dependent

By tracking and analyzing nine morphological stages as oxygen concentrations change, we identified significant differences in scaling between major morphological events. While all morphological events speed up with increasing oxygen concentrations (Figure 1C), their changes in speed are notably different. Syncytial development, as measured by the time between the appearance of the pole bud and the end of cellularization, takes proportionally less time as oxygen concentrations decrease, indicating that this stage is not slowed as much by decreasing oxygen (Figure 4A). The stages of gastrulation (end of cellularization, pole cell invagination, and amnioproctodeal invagination) are relatively uniformly affected. Germ band retraction, as measured by amnioserosa exposure, tracks subtly but inversely with syncytial development. More striking are the oxygen-dependent changes observed in head involution (clypeolabral retraction and advancing of the ventral lobe to match the clypeolabrum) and gut formation (heart-shaped midgut). While head involution takes proportionally more time as oxygen levels increase, meaning it does not slow as much as overall development in hypoxia, gut formation does the opposite. The midgut takes proportionally less time to form as oxygen levels increase, meaning it responds more strongly to increases in oxygen than overall development. This juxtaposition of behaviors leads to an inversion of when the head lobes are even versus heart-shaped midgut formation. While hypoxia leads to head involution stages finishing first, hyperoxia results in the heart-shaped midgut forming first.

**Figure 4.**
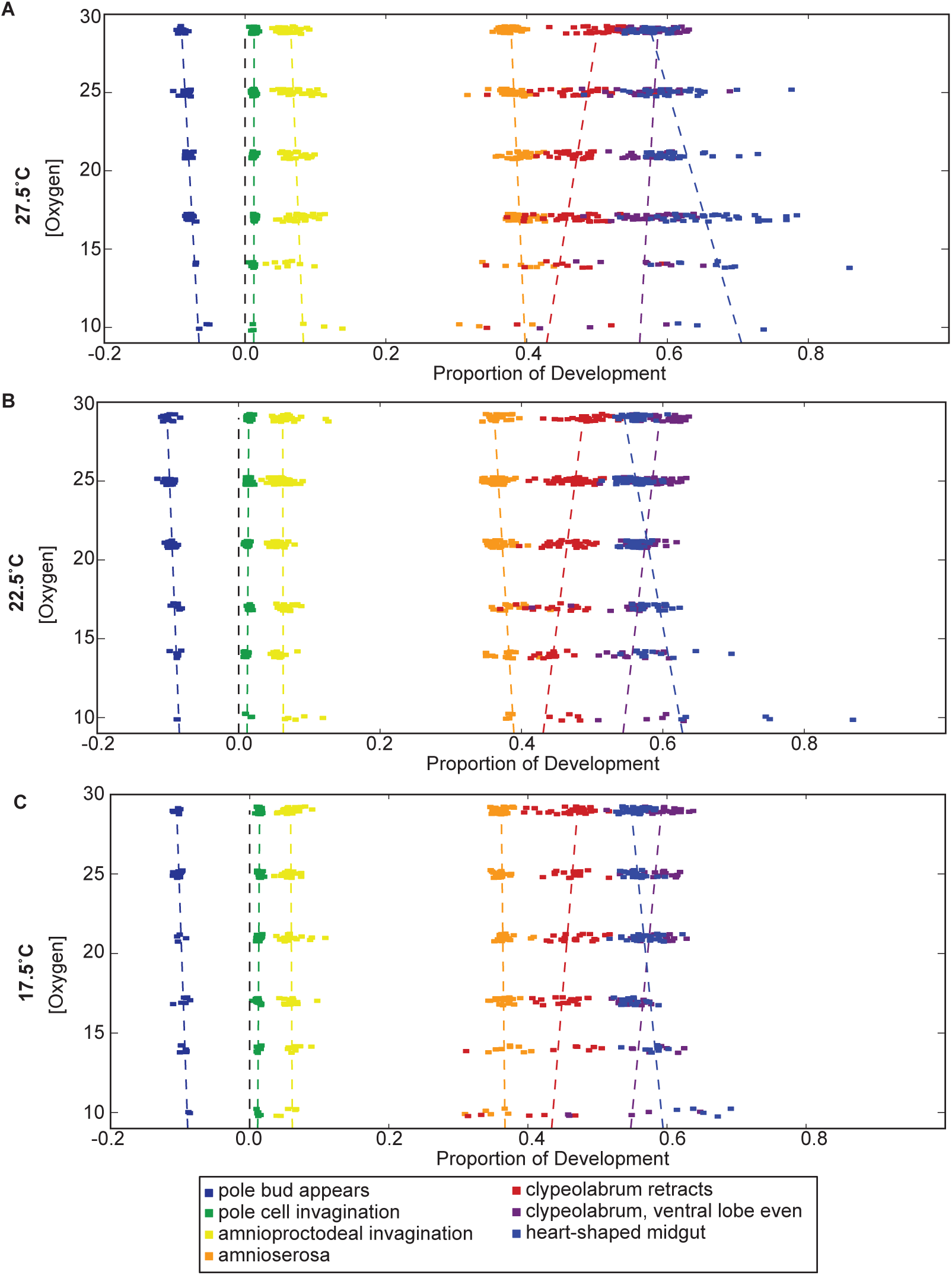
Oxygen-dependent changes vary with temperature. Gut formation and head involution are the most strikingly oxygen concentration dependent processes, but germ band retraction and syncytial development are also affected. Both syncytial development and head involution take proportionally more time as oxygen concentrations increase. Gut formation and germ band retraction, in contrast, take proportionally less time as oxygen concentrations are raised. These trends hold true across all temperatures, though the rates of change as a function of oxygen do vary. Development is normalized here between the end of cellularization and the filling of the trachea.

Surprisingly, the point of inversion varies with temperature (Figure 4). At 27.5°C, the inversion takes place at 29% oxygen, while at 17.5°C the inversion falls around 19% oxygen. This may be due to an overall shift in the oxygen response curve of heart-shaped midgut formation to proportionally later in development as temperatures fall. Supplementary Figure S2 reveals how each stage of development at each oxygen concentration changes with temperature.

## Discussion

We tracked embryogenesis at different oxygen concentrations to determine its effect on development, performing these experiments in conjunction with precise temperature control. We found that developmental rate is highly dependent on oxygen and exhibits a complex relationship with temperature. Embryos are not as robust to oxygen changes and have much less of a dynamic response than is seen with temperature. We observed significant differences in oxygen responsiveness across tissues and morphological events. These changes can be aggravated by temperature to reveal situations in which embryogenesis loses its uniform thermal scaling.

The prevalence and timing of developmental failure depend strongly on oxygen. Under hypoxia, failure is largely concentrated in mid embryogenesis at germ band retraction. Commonly the germ band fails to fully retract to expose the amnioserosa. It is possible that this stage either requires more oxygen or its complexity makes it prone to failure. A checkpoint at this stage that hypoxic embryos fail to pass may explain this phenotype. Rapid hypoxic arrests are not frequently observed under our conditions. Oxygen concentrations of 10% and 14% may fail to trigger complete hypoxic arrest in a subset of embryos yet serve to slow development enough to cause problems. Increasing oxygen levels would likely restart development, but the manner in which it restarts would depend on the stage of arrest, duration of hypoxia, and revived oxygen levels.

Hypoxia’s mid to late embryogenesis failure contrasts with high heat and high oxygen, where failure occurs during early development, during either the syncytium or early gastrulation. Under conditions with high oxygen tension, death frequently resembles, at least qualitatively, high temperature normoxia death. Failure during syncytial development commonly involves mass migration of nuclei throughout the embryo, making it difficult to distinguish the point of failure between pre-gastrulation death resulting in nuclei migration and premature gastrulation that causes death.

The syncytium responds differently to oxygen levels than other embryonic stages. Perfect scaling collapses in the syncytium at high temperatures [14], so it is not surprisingly that a difference is seen with the oxygen response as well. Interestingly, while syncytial development is less responsive to changes in oxygen than other stages across the range we tested, it is more responsive to excess heat than other stages. The difference may be aggravated by the lack of transcriptional responses available at that stage and the limited repertoire of maternally deposited genes and mRNAs. This may lead to the syncytium lacking high heat mitigations and prophylactic hypoxic responses. This implies that transcriptionally active embryos deliberately slow development either under high heat when kinetics are accelerating or under hypoxia to conserve energy.

The developmental rate response to oxygen is more subtle, yet causes more problems, across the range we tested than is seen with a moderate change in temperature. Across a 10°C differential, developmental time doubles with minimal change in viability. This is virtually invariant, regardless of oxygen concentrations. However, across a 19% change in oxygen concentration, development time experiences an absolute, rather than proportional, change of sixteen to eighteen hours. Changes in oxygen thus provide a proportionally smaller change in developmental time with enormous consequences for viability. While changes in temperature follow the Arrhenius equation, changes with oxygen appear to follow Monod’s equation. Rather than a logarithmic curve, developmental time is inversely proportional to oxygen concentration. This comparatively shallow oxygen response undermines the hypothesis that oxygen availability explains temperature-dependent changes. Changes in temperature will affect oxygen diffusion in the embryo, with a 10°C change shifting the effective oxygen concentration by *∼*4%. However, the difference in developmental time between 21% and 17% oxygen at 27.5°C is dwarfed by the dramatically larger difference between 27.5°C and 17.5°C at 21%. Therefore, basic energy metabolism is not solely responsible for the changes in developmental rates seen across temperature. Our results show that the embryo’s developmental program is robust to small changes in oxygen. This suggests some leeway in respiration efficiency; mnevertheless, there is a notable biological response.

## Materials and Methods

### Rearing and imaging of *Drosophila*

*Drosophila melanogaster*, OreR, were reared and maintained on standard fly media at 25°C. Egg-lays were performed in medium cages on 10 cm molasses plates for 1.5 hours at the temperature at which the lines were maintained after pre-clearing. Embryos were collected and dechorionated with fresh 50% bleach solution (3% hypochlorite final) for 60 seconds in preparation for imaging.

Embryos were monitored by modifying a temperature control system [14] in which an aluminum bar was embedded in an acrylic box. Both ends of the aluminum bar were external to the box and bound to Peltier heat pumps and heat sinks. A thermistor connected to the aluminum bar provided feedback to maintain the temperature using an H-bridge temperature controller (McShane Inc., 5R7-570). Embryos were glued [15] to oxygen-permeable film (lumox, Greiner Bio-one), covered with Halocarbon 700 oil (Sigma), and placed over holes drilled in the aluminum for imaging. An oxygen sensor (Grove Gas sensor (O2)) was placed in the box and connected to an external computer (Arduino-style Seeeduino V3.0 (Atmega 328P)). Finally, the box was sealed with two gas inputs and an over-pressure release. The computer utilized the oxygen sensor input and controlled two valves via NPN transistors, one connected to an oxygen tank and regulator and one connected to a nitrogen tank and regulator, to maintain specific oxygen concentrations in the box (Figure 1A).

Time-lapse imaging with bright field transmitted light was performed on a Leica M205 FA dissecting microscope with a Leica DFC310 FX camera using the Leica Advanced Imaging Software (LAS AF) platform. Greyscale images were saved from pre-cellularization to hatch. Z-stacked images were saved every two minutes (five minutes at 17.5°C). Z-stack and image analysis were conducted as previously described [14].

## Acknowledgments

We obtained flies from the Bloomington stock center. This work was supported by a Howard Hughes Medical Institute investigator award to MBE and by NIH grant HG002779 to MBE. SGK was supported by the National Institutes of Health under Ruth L. Kirschstein National Research Service Award (F32-FGM101960A) from the National Institute of General Medical Sciences. The funders had no role in study design, data collection and analysis, decision to publish, or preparation of the manuscript. We thank BCK and EDKK for their support.

## Supplementary Tables

**Table S1.** Point of Failure

| Oxygen Concentration | Temperature | Syncytial or Gastrulation Death |  | Death before Heart-shaped Midgut |  | Death before Trachea Formation |  | Failure to Hatch |  | Successful Hatch |  | Total |
| --- | --- | --- | --- | --- | --- | --- | --- | --- | --- | --- | --- | --- |
| 29% | 32.5°C | 26 | 87% | 4 | 13% | - | - | - | - | - | - | 30 |
|  | 27.5°C | - | - | 1 | 2% | 1 | 2% | 8 | 18% | 35 | 78% | 45 |
|  | 22.5°C | 13 | 28% | - | - | - | - | 2 | 4% | 31 | 67% | 46 |
|  | 17.5°C | 6 | 11% | - | - | - | - | 4 | 8% | 43 | 81% | 53 |
| 25% | 27.5°C | 3 | 8% | 1 | 3% | 3 | 8% | 2 | 6% | 27 | 75% | 36 |
|  | 22.5°C | 2 | 4% | 2 | 4% | - | - | - | - | 46 | 92% | 50 |
|  | 17.5°C | 1 | 4% | - | - | - | - | - | - | 22 | 96% | 23 |
| 21% | 27.5°C | 3 | 6% | - | - | 4 | 9% | 14 | 30% | 26 | 55% | 47 |
|  | 22.5°C | 2 | 6% | - | - | 1 | 3% | 4 | 12% | 27 | 79% | 34 |
|  | 17.5°C | 8 | 17% | 1 | 2% | 4 | 9% | 8 | 17% | 26 | 55% | 47 |
| 17% | 30.0°C | 40 | 98% | 1 | 2% | - | - | - | - | - | - | 41 |
|  | 27.5°C | 1 | 2% | - | - | 13 | 25% | 26 | 50% | 12 | 23% | 52 |
|  | 22.5°C | 6 | 13% | 2 | 4% | 12 | 25% | 10 | 21% | 18 | 38% | 48 |
|  | 17.5°C | 3 | 8% | 2 | 5% | 2 | 5% | 11 | 28% | 22 | 55% | 40 |
| 14% | 27.5°C | 19 | 38% | 17 | 34% | 6 | 12% | 8 | 16% | - | - | 50 |
|  | 22.5°C | 2 | 4% | 2 | 4% | 9 | 18% | 25 | 49% | 13 | 25% | 51 |
|  | 17.5°C | 7 | 12% | 3 | 5% | 21 | 36% | 13 | 22% | 15 | 25% | 59 |
| 10% | 27.5°C | 11 | 24% | 14 | 31% | 17 | 38% | 3 | 7% | - | - | 45 |
|  | 22.5°C | 3 | 8% | 14 | 36% | 16 | 41% | 6 | 15% | - | - | 39 |
|  | 17.5°C | 17 | 29% | 17 | 29% | 11 | 19% | 14 | 24% | - | - | 59 |

## Supplementary Figures

**Figure S1.**
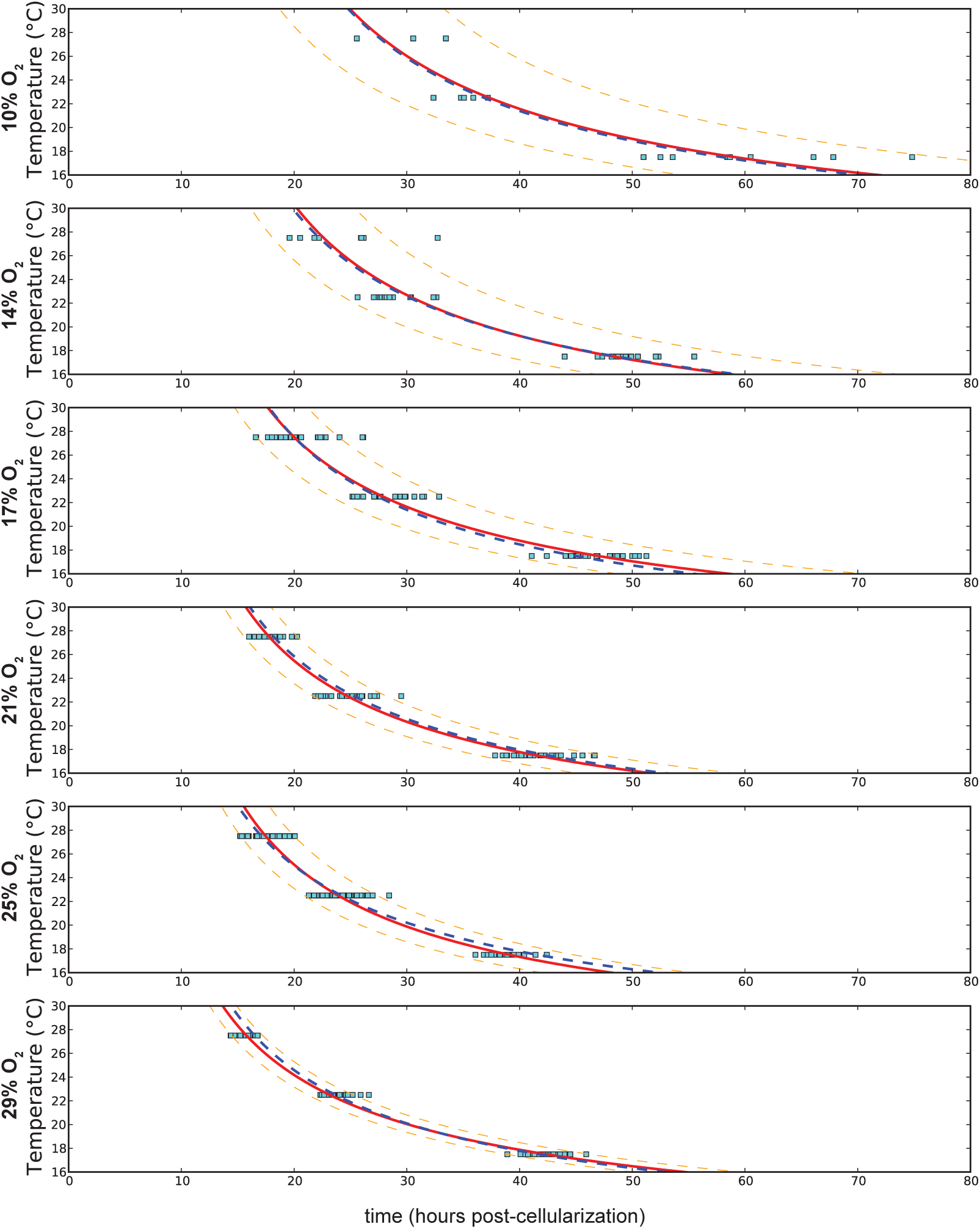
Curve fitting for each oxygen concentration as temperature changes. Curve fitting at each oxygen concentration is shown with the red line, with the 90% confidence interval delineated by the orange dashed line. The overall curve fitting is identified with the blue dashed line. Differences between different oxygen concentrations are much smaller than the range covered by temperature changes. The change in developmental time from high to low temperatures remains proportional across oxygen concentrations.

**Figure S2.**
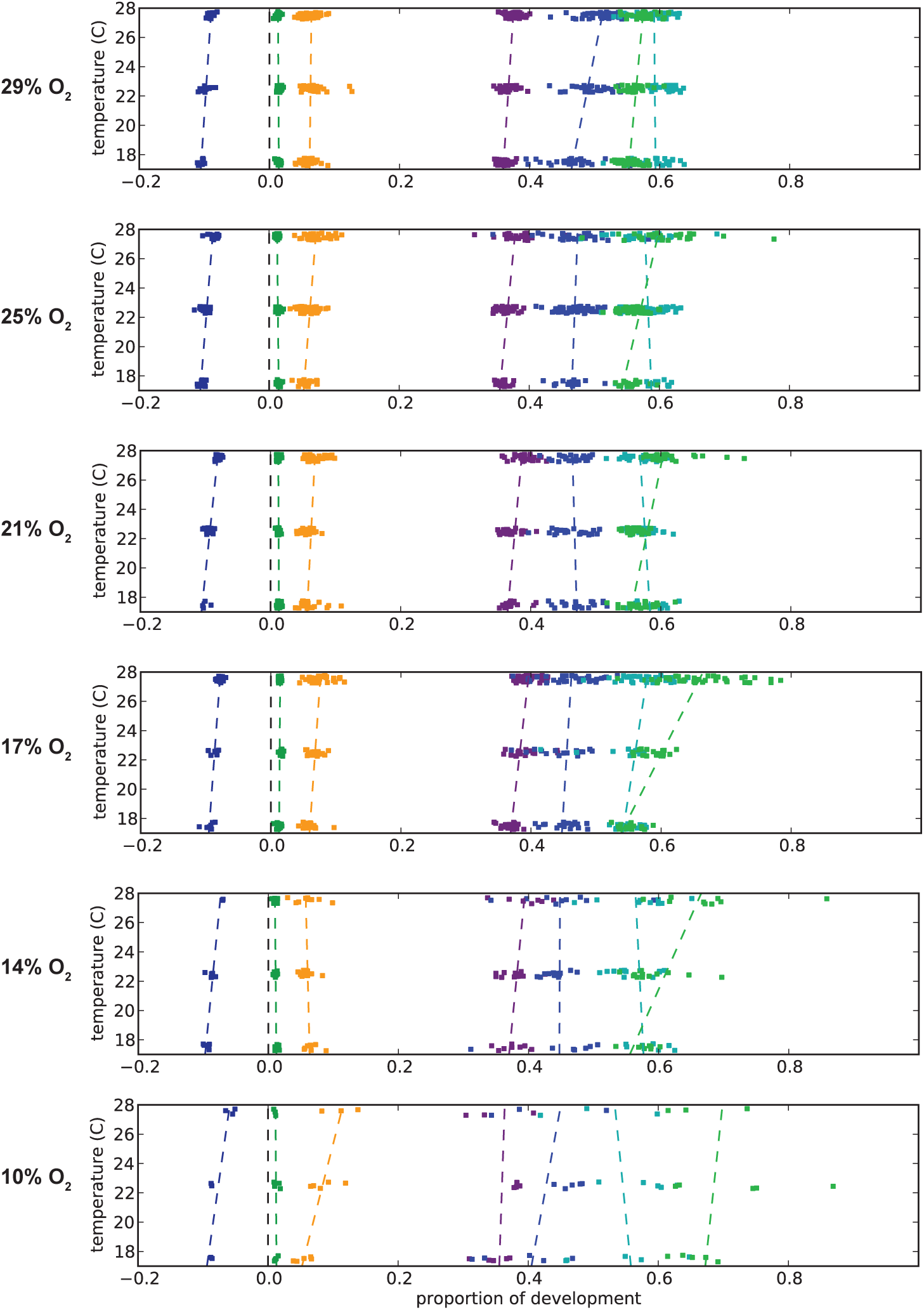
Changes in stages react differently as temperature changes. As oxygen concentrations decrease, gut-development is proportionally delayed, as can be seen with the timing of the heart-shaped midgut shifting later in development. Proportional changes in development are more severe at higher temperatures. At very low oxygen concentrations (10% oxygen), development becomes highly irregular. In many animals, development ceases during germ-band retraction at very low oxygen concentrations. Development is normalized here between the end of cellularization and the filling of the trachea.

